## Supplemental information for "Direct detection of coupled proton and electron transfers in human manganese superoxide dismutase"

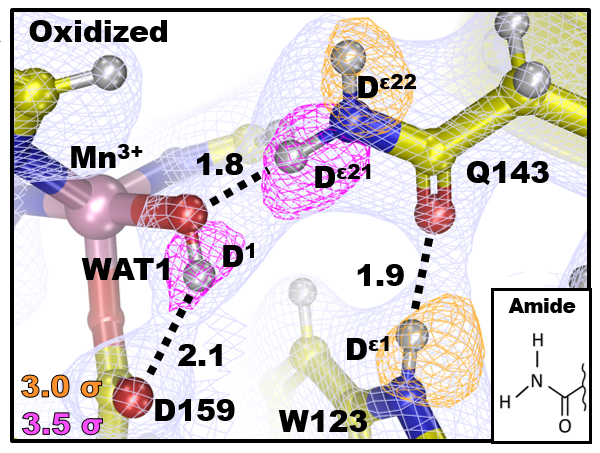

**Figure S1. Neutron structure at the active site of Mn^3+^SOD chain A.** Magenta and orange omit |F_o_|-|F_c_| difference neutron scattering length density is displayed at 3.5σ and 3.0σ, respectively, and light blue 2|F_o_|-|F_c_| neutron scattering length density is displayed at 1.0σ.

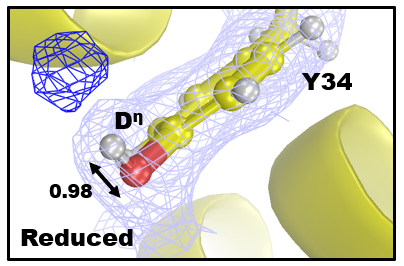

**Figure S2. Residual density for the hydroxyl group of Tyr34 in Mn^2+^SOD of chain B.** Light blue 2|F_o_|-|F_c_| neutron scattering length density displayed at 1.0 σ. Dark blue omit |F_o_|-|F_c_| difference density is displayed at 2.0 σ. Bond length in Å is given.

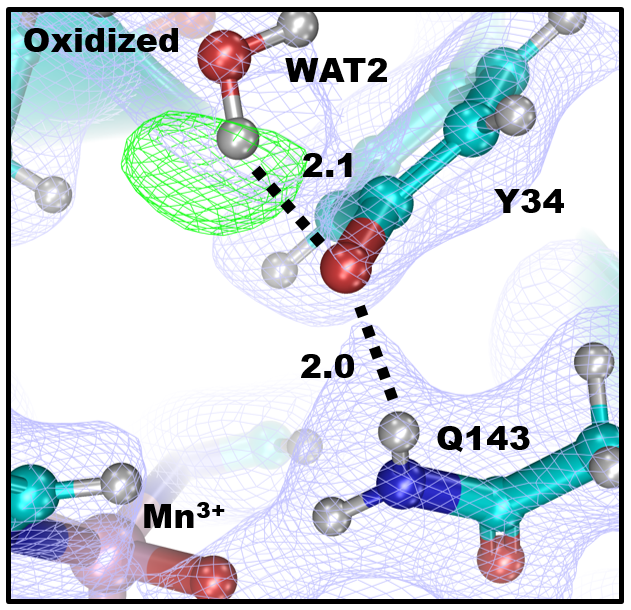

**Figure S3. Neutron structure at the active site of Mn^3+^SOD chain B.** Green omit |F_o_|-|F_c_| difference neutron scattering length density is displayed at 2.5σ and light blue 2|F_o_|-|F_c_| neutron scattering length density is displayed at 1.0σ.

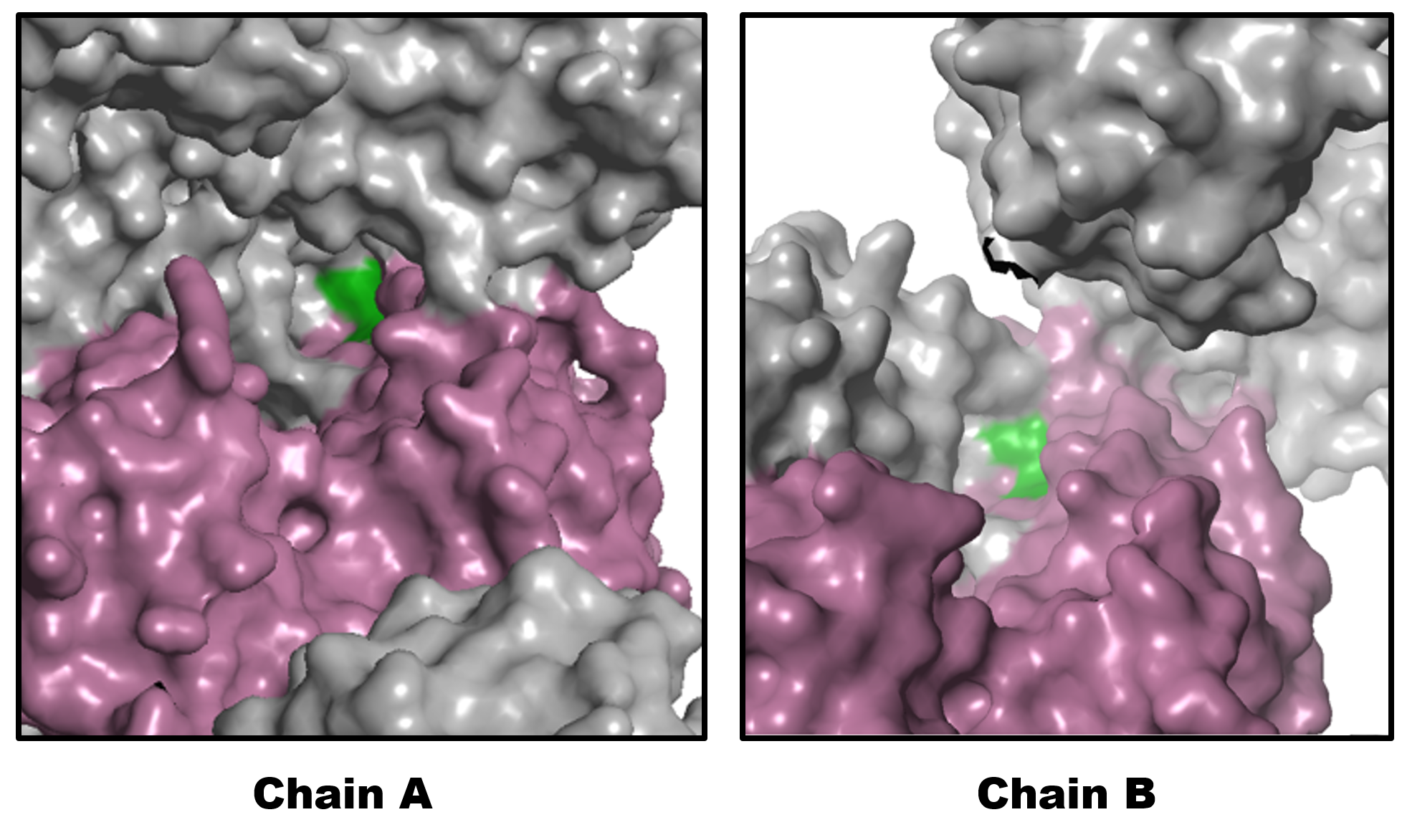

**Figure S4. Solvent accessibility differences between chains of the asymmetric AB dimer for *P*6_1_22 MnSOD.** Magenta depicts the surfaces of the chain for the asymmetric unit and green depicts the surfaces leading to the active site of the chain. Grey depicts the surfaces of symmetry generated asymmetric AB dimers.

**Table S1. Gln143 bonding character from CLPO analysis.** Numbers are calculated bond order.

|  | Five-Coordinate Mn^3+^  Y166(-)H30(ε)Y34(-) | Five-Coordinate Mn^3+^  Y166(H)H30(δ)Y34(-) | Six-Coordinate Mn^2+^  Y166(H)H30(δ)Y34(-) | Five-Coordinate Mn^2+^  Y166(H)H30(δ)Y34(H) |
| --- | --- | --- | --- | --- |
| N^ε2^-C^ε1^ | 1.36 | 1.37 | 1.56 | 1.52 |
| O^ε1^-C^ε1^ | 1.45 | 1.45 | 1.30 | 1.33 |

**Table S2. Charge and energy interactions of donor-acceptor CLPO analysis.**

| **Trp123 N^ε1^ Lone Pair → Trp123 C^ε2^-C^δ2^ π^*^-bond** | | | | |
| --- | --- | --- | --- | --- |
| State | Donor Occupancy  (e^-^) | Charge Transfer (e^-^) | Acceptor Occupancy (e^-^) | Energy Stabilization (kcal/mol) |
| Five-Coordinate Mn^2+^  Y166(H)H30(δ)Y34(H) | 1.58 | 0.15 | 0.52 | ↓ 13.52 |
| **Gln143 N^ε2^ Lone pair → WAT1 O-H σ^*^-bond** | | | | |
| State | Donor Occupancy  (e^-^) | Charge Transfer (e^-^) | Acceptor Occupancy (e^-^) | Energy Stabilization (kcal/mol) |
| Six-Coordinate Mn^2+^  Y166(H)H30(δ)Y34(-) | 1.81 | 0.14 | 0.15 | ↓ 0.46 |
| Five-Coordinate Mn^2+^  Y166(H)H30(δ)Y34(H) | 1.78 | 0.17 | 0.18 | ↓ 1.39 |

**Table S3. Percent covalence of shared hydrogen atoms in SSHBs bonds from CLPO analysis.**

|  | Five-Coordinate Mn^3+^  Y166(-)H30(ε)Y34(-) | | Five-Coordinate Mn^3+^  Y166(H)H30(δ)Y34(-) | | Six-Coordinate Mn^2+^  Y166(H)H30(δ)Y34(-) | | Five-Coordinate Mn^2+^  Y166(H)H30(δ)Y34(H) | |
| --- | --- | --- | --- | --- | --- | --- | --- | --- |
| (Gln143)N^ε2^-H-O(WAT1) | N^ε2^ | O | N^ε2^ | O | N^ε2^ | O | N^ε2^ | O |
|  | 0.92 | 0.08 | 0.92 | 0.08 | 0.29 | 0.71 | 0.36 | 0.64 |
| (Tyr166)O^η^-H-N^ε2^(His30) | O^η^ | N^ε2^ | O^η^ | N^ε2^ | O^η^ | N^ε2^ | O^η^ | N^ε2^ |
|  | 0.20 | 0.80 | 0.77 | 0.23 | 0.76 | 0.24 | 0.79 | 0.21 |
| (Tyr34)O^η^-H-O(WAT2) | O^η^ | O | O^η^ | O | O^η^ | O | O^η^ | O |
|  | 0.15 | 0.85 | 0.23 | 0.77 | 0.20 | 0.80 | 0.90 | 0.10 |
| (His30)N^δ1^-H-O(WAT2) | N^δ1^ | O | N^δ1^ | O | N^δ1^ | O | N^δ1^ | O |
|  | 0.14 | 0.86 | 0.89 | 0.11 | 0.88 | 0.12 | 0.92 | 0.08 |

**Table S4. Active Site B-factors (Å^2^) of MnSOD Neutron Structures**

|  | | **Mn^3+^SOD** | | **Mn^2+^SOD** | |
| --- | --- | --- | --- | --- | --- |
| **Molecule** | **Atom** | **A** | **B** | **A** | **B** |
| **WAT1** | **O** | **22.8** | **30.92** | **18.4** | **20.2** |
|  | **D^1^** | **27.3** | **36.99** | **21.8** | **23.9** |
|  | **D^2^** | **-** | **-** | **-** | **23.9** |
| **Q143** | **N^ε2^** | **23.1** | **21.4** | **17.8** | **27.2** |
|  | **D^ε21^** | **27.3** | **37.0** | **^b^21.8** | **-** |
|  | **D^ε22^** | **27.6** | **25.6** | **21.1** | **32.3** |
| **Y34** | **O^η^** | **20.2** | **27.2** | **25.2** | **21.9** |
| **H30** | **N^δ1^** | **24.9** | **17.8** | **29.9** | **17.1** |
|  | **D^δ1^** | **29.8** | **-** | **35.6** | **20.3** |
|  | **N^ε2^** | **24.1** | **18.7** | **23.5** | **17.7** |
| **Y166** | **O^η^** | **26.1** | **23.1** | **21.7** | **19.7** |
| **^a^H30/Y166** | **D^ε2^/D^η^** | **28.8** | **22.3** | **28.0** | **20.9** |

**^a^D atom between N^ε2^(H30) and O^η^(Y166). For Mn^3+^SOD, the atom is closest to O^η^(Y166). For Mn^2+^SOD, the atom is equidistant between N^ε2^(H30) and O^η^(Y166).**

**^b^For chain A of Mn^2+^SOD, D^ε21^(Q143) is 1.4 Å from O(WAT1) and may be partially bonded with it.**

**Table S5. Active Site Bond Lengths of MnSOD Neutron Structures**

|  | **Mn^3+^SOD** | | **Mn^2+^SOD** | |
| --- | --- | --- | --- | --- |
| **Mn Covalent Bonds (Å)** | **A** | **B** | **A** | **B** |
| Mn-N^ε2^(H26) | 2.07 | 2.07 | 2.26 | 2.10 |
| Mn-N^ε2^(H74) | 2.13 | 2.12 | 2.19 | 2.25 |
| Mn-O^ε2^(D159) | 1.95 | 1.94 | 2.44 | 2.15 |
| Mn-N^ε2^(H163) | 2.06 | 2.14 | 2.23 | 2.21 |
| Mn-O(WAT1) | 1.78 | 1.76 | 2.12 | 2.22 |
| Mn-O(OL) | **-** | **-** | 1.82 | - |

**Table S6. Data collection and refinement statistics**

| Data Collection Statistics | | | | |
| --- | --- | --- | --- | --- |
|  | Neutron | | X-ray | |
|  | **Oxidized** | **Reduced** | **Oxidized** | **Reduced** |
| PDB Code | **7KKS** | **7KKW** | **7KKU** | **7KLB** |
| Diffraction Source | MaNDi | | Rigaku FR-E SuperBright | |
| Temperature (K) | 296 | | | |
| Space group | *P*6_1_22 | | | |
| *a*, *b*, *c* (Å) | 81.30, 81.30, 241.840 | 81.33, 81.33, 242.880 | 81.14, 81.14, 241.63 | 81.13, 81.13, 242.12 |
| *α*, *β*, *γ* (°) | 90, 90, 120 | | | |
| Wavelengths (Å) | 2-4 | | 1.5418 | |
| No. of images | 8 | 8 | 180 | 280 |
| Exposure time | 48 h | 48 h | 60 s | 60 s |
| No. of unique reflections | 24556 | 21719 | 31815 | 25718 |
| Total No. of reflections | 196496 | 155817 | 248673 | 194364 |
| Resolution range (Å) | 14.64-2.20 (2.28-2.20) | 14.65-2.30 (2.38-2.30) | 50.00-2.02 (2.07-2.02) | 50.00-2.16 (2.20-2.16) |
| Multiplicity | 8.0 (6.1) | 7.2 (5.7) | 7.8 (3.5) | 7.6 (4.1) |
| I/σ(I) | 7.0 (3.40) | 6.2 (3.3) | 8.3 (2.0) | 4.8 (2.0) |
| R_merge_ | 0.284 (0.314) | 0.277 (0.294) | - | - |
| R_meas_ | - | - | 0.291 (0.610) | .459 (.683) |
| CC _1/2_ | 0.935 (0.275) | 0.943 (0.319) | 0.950 (0.730) | 0.804 (0.638) |
| R_pim_ | 0.101 (0.129) | 0.102 (0.124) | 0.082 (0.320) | 0.140 (0.326) |
| Data completeness (%) | 98.83 (98.83) | 98.94 (99.16) | 100.0 (100.0) | 97.5 (95.5) |
| Refinement Statistics | | | | |
| R_work_ | 0.2565 | 0.2493 | 0.2166 | 0.2075 |
| R_free_ | 0.2817 | 0.3021 | 0.2517 | 0.2544 |
| No. of atoms  Protein including D  Solvent  Mn | 6606  6228  376  2 | 6556  6222  332  2 | 3291  3162  127  2 | 3284  3168  98  2 |
| R.m.s. deviations  Bond lengths (Å)  Bond angles (°) | 0.093  0.78 | 0.092  0.64 | 0.002  0.47 | 0.006  0.98 |
| Average *B*-factor  Protein  Water  Mn | 33.62  30.51  25.3 | 30.57  29.19  23.53 | 28.23  30.64  17.00 | 33.82  31.91  24.30 |

**Supplementary Methods**

**Data processing and refinement.** X-ray refinement was performed by removing all non-protein entities in the starting model of 5VF9^1^, simple molecular replacement through rigid-body refinement, and subsequent restrained-positional refinement. With *COOT*^2^, the protein model was manually fit into |*F*_o_|-|*F*_c_| peaks as needed and refined first. New solvent structure and Mn atoms were manually modeled into |*F*_o_|-|*F*_c_| density. As D atoms were manually added during iterations of neutron refinement, the stereochemistry weight scale was manually adjusted due to the increased number of atoms at sensitive stereochemical positions. Loose Mn-coordination restraints were derived from the Mn^2+^SOD X-ray structure and applied to the Mn^2+^SOD neutron model, whereas the Mn^3+^SOD neutron model used restraints derived from our own DFT calculations. In both cases, the R-free value was improved with the application of these restraints.

**Computational Details.** Computations from the NWChem 6.8 software package utilized an extra-fine integration grid quadrature known to provide high precision with restricted open-shell John-Sham (ROKS) treatment^3-5^. The geometry optimizations implemented the B3LYP exchange-correlation functional dispersion corrected according to Becke and Johnson damping (DFT-D3-BJ)^6,7^. Optimizations were first performed in the gas phase utilizing an energy convergence threshold of 1 E -6 atomic units. There was no notable difference when optimizations began directly in the solution phase other than longer computational times. The nearest three water molecules found in the neutron structure counterparts, representative of the ordered solvent found at the active site, were included in the QM models in addition to the Mn-ligated solvent

**Solvation model.** The COSMO solvation model treats solvent as an implicit dielectric continuum (i.e. many solvent molecules need not be explicitly modeled in the QM system). The charge distribution of the continuum is derived using a scaled-conductor boundary condition between the cavity surface and solvent^8^. The inclusion of several explicit solvent molecules in combination with an implicit solvent model to model explicitly known hydrogen bonds increases the accuracy of energy calculations^9^. In the case of the present QM system, the explicit water molecules utilized are representative of the ordered solvent found experimentally in the neutron structures and other published X-ray structures^1^.
